# High-throughput microfluidic micropipette aspiration device to probe time-scale dependent nuclear mechanics in intact cells

**DOI:** 10.1101/641084

**Authors:** Patricia M. Davidson, Gregory R. Fedorchak, Solenne Mondésert-Deveraux, Emily S. Bell, Philipp Isermann, Denis Aubry, Rachele Allena, Jan Lammerding

## Abstract

The mechanical properties of the cell nucleus are increasingly recognized as critical in many biological processes. The deformability of the nucleus determines the ability of immune and cancer cells to migrate through tissues and across endothelial cell layers, and changes to the mechanical properties of the nucleus can serve as novel biomarkers in processes such as cancer progression and stem cell differentiation. However, current techniques to measure the viscoelastic nuclear mechanical properties are often time consuming, limited to probing one cell at a time, or require expensive, highly specialized equipment. Furthermore, many current assays do not measure time-dependent properties, which are characteristic of viscoelastic materials. Here, we present an easy-to-use microfluidic device that applies the well-established approach of micropipette aspiration, adapted to measure many cells in parallel. The device design allows rapid loading and purging of cells for measurements, and minimizes clogging by large particles or clusters of cells. Combined with a semi-automated image analysis pipeline, the microfluidic device approach enables significantly increased experimental throughput. We validated the experimental platform by comparing computational models of the fluid mechanics in the device with experimental measurements of fluid flow. In addition, we conducted experiments on cells lacking the nuclear envelope protein lamin A/C and wild-type controls, which have well-characterized nuclear mechanical properties. Fitting time-dependent nuclear deformation data to power law and different viscoelastic models revealed that loss of lamin A/C significantly altered the elastic and viscous properties of the nucleus, resulting in substantially increased nuclear deformability. Lastly, to demonstrate the versatility of the devices, we characterized the viscoelastic nuclear mechanical properties in a variety of cell lines and experimental model systems, including human skin fibroblasts from an individual with a mutation in the lamin gene associated with dilated cardiomyopathy, healthy control fibroblasts, induced pluripotent stem cells (iPSCs), and human tumor cells. Taken together, these experiments demonstrate the ability of the microfluidic device and automated image analysis platform to provide robust, high throughput measurements of nuclear mechanical properties, including time-dependent elastic and viscous behavior, in a broad range of applications.

## Introduction

The nucleus is the largest and stiffest organelle of eukaryotic cells. The mechanical properties of the nucleus are primarily determined by the nuclear lamina, a dense protein network comprised of lamins that underlies the inner nuclear membrane, and chromatin.^1–4^ Chromatin mechanics dominate the overall nuclear response for small deformations, whereas the lamina governs the nuclear response for larger deformations.^3,4^ In recent years, the mechanical properties of the nucleus have emerged as important predictors and biomarkers for numerous physiological and pathological conditions and functions, raising increased interest in probing nuclear mechanics. For example, the deformability of the nucleus determines the ability of migrating cells to pass through small openings,^5–8^ which is highly relevant during development, immune cell infiltration, and cancer metastasis, where cells move through tight interstitial spaces and enter and exit blood vessels through openings only a few micrometre in diameter.^9^ In stem cell applications, the morphology and mechanical properties of the nucleus can serve as label-free biomarkers for differentiation,^10–12^ reflecting characteristic changes in the composition of the nuclear envelope and chromatin organization during differentiation.^10,13,14^ Lastly, mutations in the genes encoding lamins give rise to a large family of inheritable disorders termed laminopathies, which are often characterized by reduced nuclear stability.^15^

The mechanical properties of cells and their nuclei are assessed using a range of techniques. Nuclear deformation can be observed by stretching cells cultured on flexible membranes and used to infer the mechanical properties of the nucleus, including the contribution of specific nuclear envelope proteins.^16–19^ However, this technique relies on nucleo-cytoskeletal connections to transmit forces to the nucleus, which may be affected by mutations in nuclear lamins,^20^ and stretching cells requires strong adhesion to the substrate. The latter fact limits the type of cells that can be studied, and can result in bias towards sub-populations of strongly adherent cells.^19^ Single cell techniques, such as atomic force microscopy (AFM), nuclear stretching between two micropipettes,^4^ and magnetic bead microrheology,^21^ apply precisely controlled forces and measure the induced deformation, thus providing detailed information on nuclear mechanical properties. However, these techniques are time-consuming, technically challenging, and often require expensive equipment and training.

Micropipette aspiration remains one of the gold standards and most commonly used tools to study nuclear mechanics^22–24^ and provides important information on the viscoelastic behaviour of the nucleus over different time scales.^13,25^ Micropipette aspiration has been used to study a wide variety of phenomena, including the mechanical properties of the nucleus^2,25^, the exclusion of nucleoplasm from chromatin,^26^ and chromatin stretching^27^ during nuclear deformation. However, micropipette aspiration is traditionally limited to a single cell at a time and performed with custom-pulled glass pipettes, which often vary in shape and diameter. In contrast, microfluidic devices enable high-throughput measurements of nuclear and cellular mechanics with precisely defined geometries.^28–30^ Some microfluidic devices measure the stiffness of cells based on their transit time when perfused through narrow constrictions^31–34^ or mimic micropipette aspiration,^35^ but these approaches are often hampered by clogging due to particles, large cell aggregates, or cell adhesion in the constrictions. This problem can be alleviated in devices that use fluid shear stress to deform the cells rather than constrictions,^36^ but the deformations achieved in these devices do not recapitulate the extensive deformations that can be achieved using physical barriers. Furthermore, in many of the current microfluidic perfusion assays, nuclear deformation is measured for only fractions of a second, making it difficult to observe viscoelastic responses with longer time-scales.

To overcome these challenges, we have developed an easy-to-use microfluidic device to measure time-dependent nuclear mechanical properties in a high-throughput manner. The device enables robust measurements of many cells in parallel, with easy loading and removal of cells from the aspiration sites, and requires minimal specialized equipment. Combined with a custom-developed automated image analysis MATLAB program to further accelerate the analysis and to provide consistent measurements, this experimental platform enables analysis of 100’s of cells per hour, representing a 10-to 40-fold improvement over conventional manual micropipette aspiration.^37^ We demonstrate the device’s utility to quantify time-dependent nuclear and cell mechanics on a single-cell level, in a high throughput manner, in a broad range of applications and cell types.

## Materials and Methods

### Cells used for experiments

Mouse embryonic fibroblasts (MEFs) with homozygous deletion of the *Lmna* gene, which encodes lamins A/C, along with wild-type littermate controls, were generously provided by Dr. Colin Stewart.^38^ Wild-type MEFs were stably modified with lentiviral vectors to express mNeonGreen-Histone 2B,^39^ as described previously.^40^ HT1080 cells were purchased from the DSMZ Braunschweig, Germany, and stably modified with lentiviral vectors to express the nuclear rupture reporter NLS-GFP, as described previously.^41^ Induced pluripotent stem cells (iPSC) and healthy human skin fibroblasts were generously provided by Elisa di Pasquale and Gianluigi Condorelli (Humanitas Clinical and Research Center, Italy).^42^ MDA-MB-231 cells were obtained from the American Type Culture Collection (ATCC). MEF, HT-1080, MDA-MB-231, and human fibroblast cells were maintained in Dulbecco’s Modified Eagle Medium (DMEM) supplemented with 10 % (v/v) fetal bovine serum and 1 % (v/v) penicillin/streptomycin. iPSCs were maintained on matrigel-coated dishes in mTeSR medium (Stem Cell Technologies), prepared according to manufacturer’s instruction. The dishes were prepared by diluting 50 µl matrigel (BD 354277) in 1 ml of mTeSR and incubating in 35 mm plastic petri dishes overnight at 4°C.

### Design and microfabrication of the microfluidic devices

The mask and wafers were produced in the Cornell NanoScale Science and Technology Facility (CNF). The masks were fabricated using a Heidelberg DWL 2000 Mask Writer. Since the device contains features with different heights (5 µm for the micropipette channels and 10 µm for larger perfusion channels), two SU8 photolithography steps were used. A first 5-μm tall layer consisting of only the micropipettes channels was created by spinning SU-8 2005 to the correct thickness and exposing through the photomask using a GCA Autostep 200 DSW i-line Wafer Stepper, which allows precise realignment of the mask and wafer within 1 μm when using masks for the different SU-8 layers. The wafer was baked at 95°C for 30 minutes, cooled down and developed in SU-8 developer. A second layer of SU-8 2007 was spun to a thickness of 10 μm, and the larger device features were exposed on the stepper. The wafers were subsequently baked, developed following standard photolithography procedures,^40^ and coated with trichloro(1H, 1H, 2H, 2H-perfluorooctyl)silane to facilitate demolding. PDMS replicas of the devices were cast using Sylgard 184 (Dow Corning), mixing in a 10:1 ratio and baking for two hours at 65°C. To minimize wear to the original wafer, the first PDMS cast was used to create a plastic mold from which all subsequent PDMS replicas were made, following a previously published protocol.^43^ PDMS replicas were cut into individual devices and holes for perfusion were cut into the PDMS using a small (0.75 or 1.2 mm) biopsy punch to introduce tubing. The final PDMS devices were then mounted on glass slides using a plasma cleaner (Harrick Plasma) as described previously.^5,40^

### Experimental acquisition

Immediately after plasma treatment, the PDMS devices were filled with 20 mg/ml bovine serum albumin (BSA) and 0.2% (v/v) fetal bovine serum (FBS) in phosphate buffered saline (PBS) for 10 minutes to passivate the device. The same PBS solution was used as perfusion buffer and to create a cell suspension. The cell suspension (5 million cells/ml) was prepared in the PBS solution and kept on ice. Cell nuclei were stained by adding an aliquot of Hoechst 33342 at a dilution of 1:1000 to the cell suspension for a final concentration of 10 µg/ml and incubated on ice for ten minutes before being used for experiments. The vial with the cell suspension was connected via Tygon S3 E-3603 tubing (VWR, inner diameter 1/32"; outer diameter 3/32”) to the cell entry port of the microfluidic device; a vial with cell free PBS solution (perfusion buffer) was connected to the buffer port. Additional tubing was connected to the outlet port (*P*_atm_ in Figure 1A) and drained into a small collection tube. The pressure applied to the vials with the cell suspension and the perfusion buffer was adjusted using an MCFS-EZ pressure controller (Fluigent) to regulate cell/buffer perfusion into the device. For the experiments, a pressure of 7.0 kPa was applied to the cell suspension and 1.4 kPa to the buffer solution. The outlet port tubing was open to atmospheric pressure.

**Figure 1:**
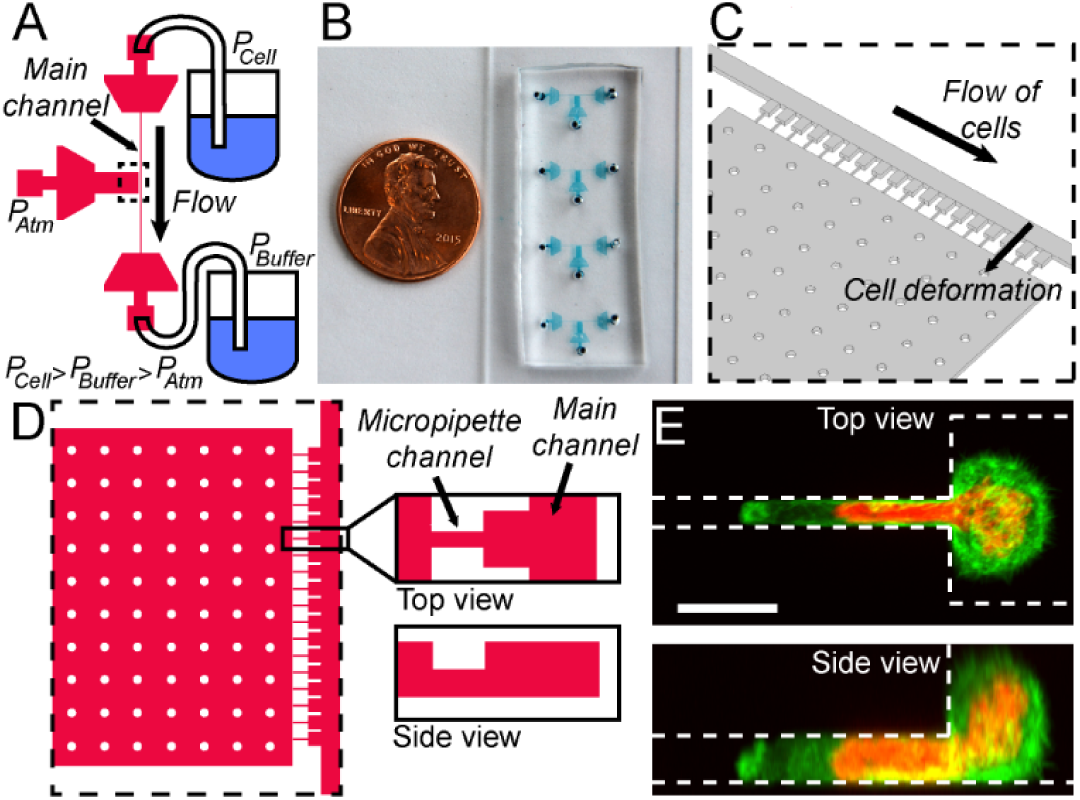
Overview of the micropipette devices. (**A**) Schematic overview of the device and the different pressures applied to the three ports. The dashed rectangle indicates the region shown as close-up in panels C and D. (**B**) Photograph of the actual devices in a typical experimental setup, with four devices mounted on a glass coverslip, allowing the measurement of four different cell types or replicates in rapid succession. A US 1 cent coin serves as reference for size. (**C**) Schematic 3-D close-up of the micropipette channels and the main channel, corresponding to the area outlined with a dashed line in panel A. (**D**) Schematic close-up of the device region with the individual pipettes channels, viewed from the top (left) and side (right). The side-view shows that the pipette channels have a lower height (5 µm) than the rest of the device (10 µm). (**E**) Representative image of cells expressing fluorescently labelled histones (red) to reveal the nucleus, and a fluorescent actin marker (LifeAct-GFP, green) to delineate the cytoplasm, entering the micropipette channel. (Scale bar: 10 µm)

Brightfield and fluorescence images of cells in the micropipette channels were acquired every 5 seconds using a 20×/NA 0.8 air objective and ORCA Flash 4.0 V2 Deep Cooling sCMOS (Hamamatsu) or alternatively CoolSNAP KINO CCD (Photometrics) digital camera to record nuclear deformation. At the start of each acquisition, cells present in the device were ejected from the micropipette channels, allowing new cells to enter the cell pockets and micropipette channels. To eject cells, pressure was applied to the outlet port with a syringe or pipette inserted in the tubing, causing transient reversal of the flow in the micropipette channels. As the pressure is greater at the cell port than the buffer port, the ejected cells were swept away from the vicinity of the micropipette channels towards the buffer port. After these cells had been removed (as observed through the microscope), the pressure at the outlet port was released, allowing new cells to enter the cell pockets and micropipette channels. The next round of data acquisition was then performed with these cells. By commencing the image acquisition before ejecting the cells, we ensured that all stages of cell and nuclear deformation were captured in the image sequences. The above procedure was repeated several times to capture data for a large number of cells at each experimental condition.

### Modelling and experimental validation of fluid dynamics in the microfluidic devices

To determine the pressure exerted on the cells during nuclear deformation in the micro-channels, and because physical measurements inside the device are not feasible, we computationally modelled the pressure distribution inside the devices. Using the finite elements modelling software COMSOL Multiphysics 5.2, we designed a three-dimensional (3D) model that reproduced the geometry of the device. The fluid flow in the device was considered as laminar flow following the Navier-Stokes equation:

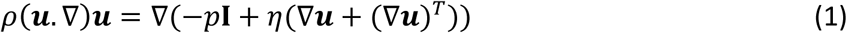

in which *ρ* is the volumic mass, ***u*** is the velocity, *p* is the pressure, *I* is the identity matrix and *η* is the dynamic viscosity of the fluid. The operator *T*, indicates the transpose operation on a tensor.

The hydrodynamic resistance of a tubular channel with laminar flow scales with the length of the channel and the inverse of the channel radius to the fourth power. Since the cross-sectional area of the tubing connecting the pressure controller to the device is orders of magnitude larger than the cross-sectional area of the channels in the microfluidic device, the hydrodynamic resistance of the microfluidic device is much greater than that of the connecting tubing. The pressure drop across the tubing outside of the microfluidic devices was therefore considered negligible relative to the pressure drop inside the device. The boundary conditions of the model were thus set to the pressure values applied to each solution in the device (*P*_Cell_ = 7 kPa; *P*_Buffer_ = 1.4 kPa). From this simulation, we computed the pressure distribution and the corresponding fluid flow profile in the device. The simulated velocity field was averaged over surfaces located above the centre of each pocket, to remove any effects due to variation in the geometry.

To validate our computational model, we experimentally determined the flow rates from the streaks created by fluorescent beads (1.9 µm diameter) over a 3 ms exposure time. The length of the streaks was measured using ImageJ (National Institutes of Health, https://imagej.nih.gov/ij/). To minimize the effect of bead interactions with the walls, we analysed only beads in the centre of the channel. Given the small dimensions of the microfluidic channels, we calculated the effect of the beads on the effective viscosity of the fluid, using the work of Heinen et al.^44^ and Einstein’s formula:

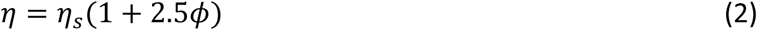

in which *η*_*s*_ represents the dynamic viscosity of the fluid alone, and *ϕ* is the volume fraction of beads in the fluid. In our experiments, we used a 0.01% vol/vol suspension of beads with 1.9 µm diameter (Thermo-Fisher, Fluoro-Max G0200) in PBS solution with 20 mg/ml BSA. The viscosity for PBS containing 20 mg/ml BSA is ⍰_s_ = 1.12 mPa.s.^44^ Using the above equation, the dynamic viscosity of the bead/PBS suspension was determined from equation (2) to be ⍰ = 1.148 mPa.s. The flow rates in the channels were then computed from equation (1) using the bead velocity, pressure, and the viscosity of the bead solution.

### Automated analysis of nuclear deformability measurements

A custom-written MATLAB program (available at: https://github.com/Lammerding/MATLAB-micropipette_analysis) was used to compute nuclear deformation into the microfluidic micropipette channels with only minimal user intervention. The MATLAB script converts time-lapse micropipette aspiration movies obtained using ZEN software (Zeiss) into multidimensional TIF stacks, separated according to color channels. The program can be readily adapted to import time-lapse sequences in other formats. The program automatically aligns the image sequence to a mask of the microfluidic device features to correct the images for rotational error, segment the individual microfluidic pockets, and determine the location of the micropipette channel entrances. The user can make manual fine adjustments to the micropipette entrance line at any time using the arrow keys in the program interface. The program then thresholds the blue colour channel, which corresponds to the blue fluorescence from the DNA-binding Hoechst 33342 dye, to provide a trace of the nucleus during deformation. The threshold for the nuclear segmentation is based on a manual graphical user interface that provides a preview of the segmentation. To account for the heterogeneity in the Hoechst signal across different nuclei, the user selects a binary threshold value for each pocket from a histogram of pixel count versus intensity. After applying erosion and dilation processing to smooth the outlines of each thresholded nucleus, the program employs the MATLAB’s *regionprops* function to track the nucleus’ leading edge inside the micropipette and calculate the distance between the leading edge of the nucleus and the micropipette channel entrance for each frame. The program allows for visual inspection of the nuclear protrusion length analysis. After analysing all nuclei, the program exports the final matrix of nuclear protrusion values over time into a Microsoft Excel-compatible file, where rows correspond to the pocket number and columns to each image frame/time point. Empty pockets register as zeroes. Likewise, once a nucleus deforms past the end of the micropipette channel, it also registers as zero since the protrusion length is no longer measureable. For cells with highly deformable nuclei, multiple cells may sequentially enter and pass through a given micropipette channel during a single acquisition sequence. These cells are recorded as separate events. An additional MATLAB script, available upon request, was used to transpose the protrusion length versus time data to make it suitable for multilevel model analysis using JMP software.

### Fitting the deformation data to models

The data obtained in the deformation experiments were fit to a number of viscoelastic models using the solver function in Microsoft Excel. Briefly, the function corresponding to the model studied was determined and approximate values for the variables were chosen as starting values. A computed value of the protrusion length was then obtained for each given deformation time, based on the function and variables. Each of these calculated values was subtracted from the value of the protrusion length obtained experimentally at each time point. This residual value was squared and the sum of squares for all time points was used as an indicator of goodness-of-fit. The solver function in Microsoft Excel was used to minimize the sum of the squared residuals by varying the variables within each model.

Each data set was modelled using six separate functions. We tested two functions for the power law model: *y* = *A* * *t* ^⍰^ and *y* = *A* * *t* ^⍰^+ *c*. We tested four functions for the modified spring-and-dashpot model: the Kelvin–Voigt model (spring and dashpot in parallel) *y* = *A* * (1 – exp(*B* * *t*)), the linear model (a spring followed by a spring and dashpot in parallel) *y* = *A* – *B*\*(1 – exp(*C* * *t*)), a Jeffreys model (a dashpot followed by a spring and dashpot in parallel) *y* = *A* * (1 – exp(*B* * *t*)) + *C* * *t*, and a Burgers model (a spring and dashpot in series followed by a spring and dashpot in parallel) *y* = *A* – *B* * (1 – exp(*C* * *t*)) + *D* * *t*. In the results section we report the second power law model and the Jeffreys model, which both showed significant improvements over more simple models. The Burgers model did not greatly improve the sum of the residuals, and thus we chose the Jeffreys model. The viscosity and elastic modulus were derived from these variables as detailed in the Supplementary information. We calculated and report the coefficient of determination (*R*^2^) value for each model and cell type.

### Statistical analysis

Statistical analysis was performed using Microsoft Excel and Igor Pro. We determined *p* values in student *t*-tests using the TTEST function in Excel. Igor Pro was used to obtain the confidence interval (one standard deviation) on the variables obtained from the fit of the data to the various models. Standard error propagation calculations were performed to obtain error values on the spring constants, elastic moduli, and viscosities, estimating that the error on the pressure is 0.3 kPa, and the error on the width and height of the micropipette channels is 0.5 µm. In all figures, error bars represent the standard error of the mean unless indicated otherwise. All data are based on at least two independent experiments.

## Results and Discussion

### Design of the microfluidic devices

The device consists of a series of 18 pockets with small micropipette channels, abutting a larger main channel used to perfuse cells into the device and the individual pockets (Figure 1A-D). The pockets are 20 µm wide and 10 µm tall, thus large enough to hold only a single cell. The micropipette channels are 3 µm wide and 5 µm tall, similar in size to micropipettes in conventional micropipette aspiration assays for probing nuclear mechanics, in which pulled glass pipettes with 3-5 µm inner diameter are used.^2,25,45^ The micropipette channels connect to a large chamber at atmospheric pressure (*P*_atm_). The cells are introduced into the device at the cell port under a pressure (*P*_Cell_) that is higher than the pressure at the buffer port (*P*_Buffer_), ensuring that the cells flow along the main channel (Figure 1A, C and D.) The two pressure inlets allow precise control of the velocity of the perfusion of the cells through the devices and the pressure applied on the cells in the pockets and micropipette channels. Microfluidic filters at each port, consisting of arrays of pillars, prevent large clusters of cells or dust to enter the main channel. As cells perfuse through the device, single cells flow into empty pockets and block the entrance of the micropipette channels, thereby preventing additional cells from settling into the same pocket. Cells located in the pocket then deform into the micropipette channels as they are subjected to the pressure difference between the main channel and atmospheric pressure. The large cell nucleus fills the entire cross-section of the micropipette channel (Figure 1E). The externally applied pressure is kept constant and the nucleus gradually enters the micropipette channel, closely resembling the creep behaviour observed in conventional micropipette aspiration assay.^22,37^ The deformation of the nucleus over time is recorded by time-lapse microscopy and used to infer the mechanical properties of the nucleus. Our micropipette dimensions are optimized for fibroblasts, myoblasts, and most cancer cells. The design can readily be adapted for smaller, more deformable cells (such as immune cells) if needed.

### Automated image analysis

To measure nuclear deformations into the array of micropipettes in a quick and highly consistent manner, we developed a semi-automated MATLAB image analysis platform that requires only minimal user input (Figure 2). After initial image processing, a mask alignment step corrects the images for rotational error, segments the individual pockets, and determines the micropipette entrance (Figure 2B, vertical yellow line). To account for the heterogeneity in the nuclear fluorescence signal (e.g., DNA fluorescently labelled with Hoechst 33342), the user selects a binary threshold value for each pocket from a histogram of pixel count versus pixel intensity (Figure 2B, middle panel). Following additional erosion and dilation processing to smooth the segmented nuclei, the program tracks the leading edge of each nucleus (Figure 2B, red vertical line) and calculates the distance aspirated into the micropipette channel (i.e., the protrusion length) for each frame (Figure 2C). The program allows visual inspection of the nuclear protrusion length in each pocket before proceeding. The program exports the final matrix of nuclear protrusion values over time for each pocket as an Excel-compatible file for subsequent statistical analysis or curve fitting.

**Figure 2:**
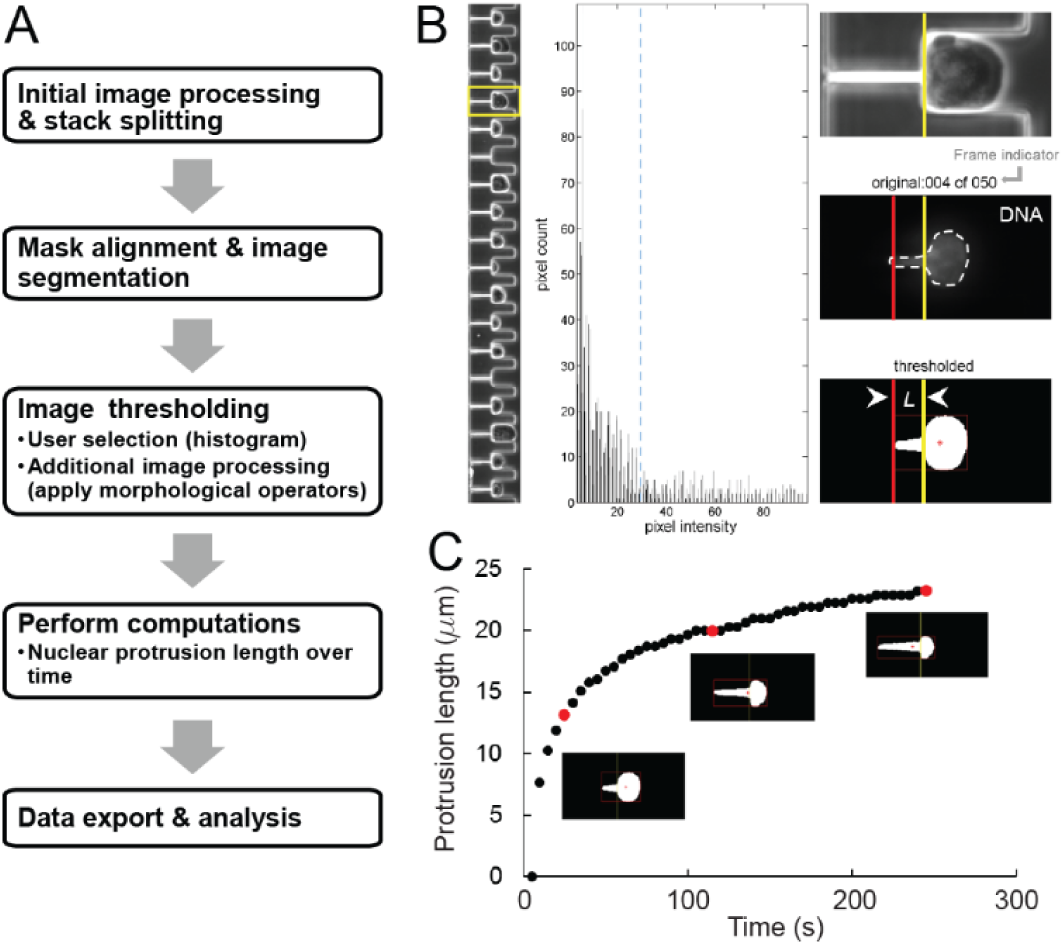
Custom-designed MATLAB software enables rapid analysis of nuclear deformability. (**A**) Schematic overview of the image analysis pipeline. The MATLAB program converts time-lapse micropipette aspiration movies into multidimensional image stacks and separates them by colour channel. The user aligns a mask to one of the image frames to segment the 18 pockets, enabling individual examination of each cell nucleus. (**B**) A graphical user interface ensures accurate measurement of the nuclear deformations within each pipette. The yellow box (left panel, fourth pocket) indicates the selected cell and corresponding nucleus, as visualized using Hoechst 33342 dye, which fluorescently labels DNA. The user sets a binary threshold value (blue dotted line) by clicking within the middle panel, a 60-bin histogram of image intensity values. Clicking the left mouse button previews the threshold by playing through the image sequence (right panel) at a user-specified sampling rate (every n^th^ frame). Additional erosion and dilation processing steps smooth boundaries and remove spurious pixels within the thresholded image. The program computes the nuclear protrusion length at each frame by drawing a bounding box around the thresholded nucleus (red box) and then computing the distance between the left edge (red vertical line) and the start of the micropipette channel (yellow vertical line). Once the thresholded image sequence (right panel, bottom) accurately depicts the original (right panel, middle), right clicking the mouse button saves the protrusion length values and proceeds to the next pocket. The values are exported to an Excel file where they can be plotted and analysed. (**C**) A plot of the nuclear protrusion length over time for a given cell, with the red data points corresponding to the thresholded nuclei in the frames shown below.

### Characterization of fluid dynamics and pressure gradients with the microfluidic device

The velocity of the cells moving along the main channel depends on the difference between the applied pressures, *P*_Cell_ and *P*_Buffer_. The larger the pressure gradient, the faster the cells will move through the device, ensuring rapid filling of available pockets. The pressure difference across the micropipette channels drives the cell and nuclear deformation. This pressure gradient is determined by the pressure in the main channel in front of the pipette (which depends on *P*_Cell_ and *P*_Buffer_) and the atmospheric pressure, *P*_atm_, at the other end of the micropipette channel. The deformation rates and flow velocities are thus readily tunable by varying the pressures applied to the cell port (*P*_Cell_) and the buffer port (*P*_Buffer_). To determine the pressure distribution within the device in more detail, including potential differences in the pressure exerted across the 18 parallel micropipette channels, we performed computational modelling of the fluid dynamics and pressure drop across the microfluidic device and then compared these model predictions with experimental measurements. We modelled two cases: one in which the micropipette channels are unfilled (“open”), and one in which the channels are blocked (“closed”). Typical experimental conditions during nuclear deformation measurements correspond to the “closed” scenario, as all of the micropipette channels are rapidly filled with cells that occupy the entire cross-section of the channels (Figure 1E) and thereby block fluid flow across the microchannels, in agreement with previous work.^46^ In the closed case, the model predicts a linear decrease in pressure across the micropipette channels (Figure 3A, B), with the cells in the micropipette channels exposed to pressures between 3.8 and 4.4 kPa, corresponding to a difference of approximately 15% between the first and the last micropipette channel. In the case of the open micropipette channels, the model predicts a pressure drop across the main channel at the pipettes that decreases rapidly. In this case, the pressure difference from the first to the last micro-pipette channel decreases from 2.4 to 1.7 kPa (Supplemental Figure 1), a difference of >40%, which would imply a large variation from one micropipette channel to the next. In both the “open” and “closed” cases, the model indicates that the pressure drop across the filters at the ports is negligible compared to the pressure drop along the main channel (Figure 3A, B; large triangular shaped areas at each of the three outlets).

**Figure 3:**
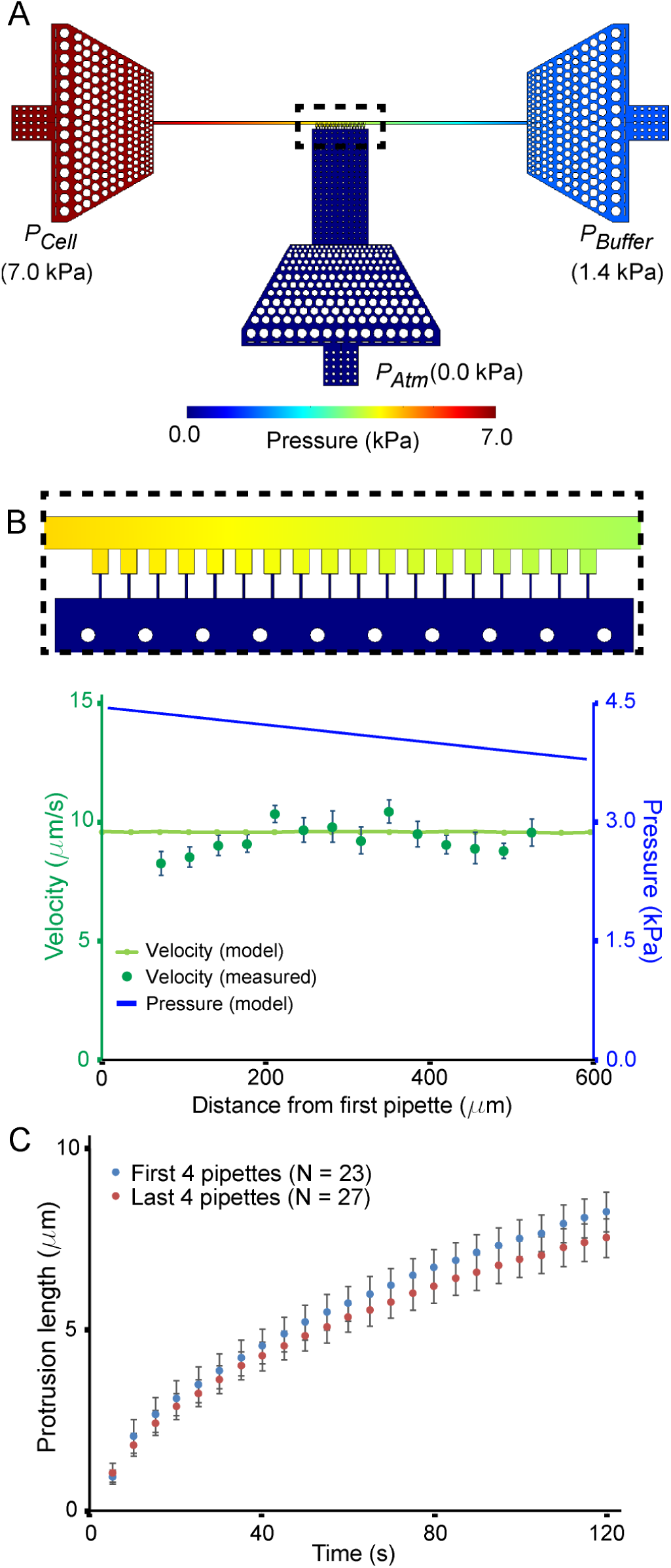
Modelling of the pressure distribution across the device. (**A**) Pressure distribution obtained from 3-D computational model in the condition in which the micropipette channels are closed, corresponding to experimental conditions in which all the channels are blocked by cells. (**B**) Comparison of the model predictions for the pressure distribution and resulting fluid velocity distribution in the main channel with experimental measurements. The velocity (light green line) determined from the pressure gradient (top figure and blue curve) was compared to the flow velocity determined from fluorescent beads (dark green points). (**C**) Deformation of wild-type MEFs in the first four micropipette channels (blue) compared to the last four micropipette channels (red). The differences between the first four and the last four channels is not statistically significant, consistent with the predictions of the models. Similar results obtained from independent experiments with another cell line are included in Supp. Fig. 2A.

### Experimental validation of the computational model

The small dimensions of the microfluidic device prohibit direct pressure measurements within the device. We therefore used experimental measurements of the fluid flow to infer the local pressure variation within the device. For these experiments, we perfused fluorescent beads through the microfluidic devices and determined the flow velocity inside the devices by quantifying the local velocity of the fluorescent beads. Measurements were obtained before and after the beads had clogged the microchannels, simulating the “open” and “closed” configurations, respectively. The experimental velocity measurements closely matched the predicted velocity from our computational model in the corresponding configurations (Figure 3B and Supp. Fig. 1). During actual micropipette aspiration experiments, all of the microchannels are simultaneously filled with cells, and thus experimental conditions resemble the “closed” case, resulting in a small, linear pressure drop along the length of the main channel. We tested whether the predicted small pressure difference between pipettes can affect the experimental readings depending on the position of the specific micropipette channel by performing experiments with mouse embryo fibroblast (MEF) cells and human breast cancer cells. The experiments did not reveal any statistically significant difference between the extent of nuclear deformation in the first 4 channels of the devices compared to the last four channels for either of the cell lines (Figure 3C; Suppl. Fig. 2A), indicating that the small drop in pressure along the main channel predicted by numerical simulations (Figure 3B) is negligible compared to the cell-to-cell variability of the experiment. If desired, the device design could be readily adapted to reduce further the pressure gradient across the section of the main channel containing the cell pockets, for example, by lengthening the other sections of the main channels, or altering its cross-section.

### Device validation in cells with known nuclear mechanical properties

To validate our microfluidic micropipette devices, we measured the nuclear mechanical properties of lamin A/C-deficient (*Lmna*^−/–^) and wild-type (*Lmna*^+/+^) MEFs, which have been extensively characterized by micropipette aspiration^5^ and nuclear strain experiments^1,47^. Consistent with previous studies, we found that lamin A/C-deficient MEFs had significantly more deformable nuclei than wild-type MEFs, as evidenced by the substantially more rapid deformation into the micropipette channels (Figure 4). Lamin A/C-deficient cells exhibited nuclear deformations 2.17 ± 0.02 times larger than wild-type controls, which is similar to the 2.05-fold increase in nuclear deformation observed in the same cell lines using substrate strain experiments,^1^ and the 2.2-fold increase reported in a previous study comparing lung epithelial cells depleted for lamin A/C to non-depleted controls.^48^ For a more detailed analysis of the mechanical properties of these two cell types, we compared the time-dependent nuclear deformation into the micropipette channels using two alternative approaches. In the first approach, we modeled nuclear deformation into the micropipette channels under a constant pressure (‘creep’) using a power law proposed by Dahl and colleagues.^25^ In this model, the nuclear protrusion length increases as a function of time to the power of an exponent, *α*, and the prefactor, *A*; the constant C accounts for uncertainty in the exact timing when the nucleus entered the channel (*t* = 0).

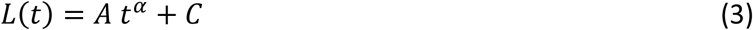

For viscoelastic materials, the exponent *α* is in the range of 0 to 1, and indicates whether the material behaviors more elastic(*α* closer to 0) or more viscous (*α* closer to 1).^25^ In our experiments (Figure 4B), lamin A/C-deficient and wild-type cells both fit power laws with similar exponents (*α* = 0.41 ± 0.01 and *α* = 0.37 ± 0.01 for wild-type cells and lamin A/C-deficient cells, respectively). This value is comparable to the one found by Dahl et al.^25^ (*α* = 0.3) for human adenocarcinoma-derived epithelial-like cells (TC7), and in agreement with a later study by the same group that found that reducing lamin A/C levels does not significantly affect the power law exponent for time-scales exceeding 10 seconds (*α* = 0.20 for wild-type and 0.24 for lamin A/C-depleted lung epithelial cells).^48^ Taken together, our data indicate that the microfluidic devices produce results consistent with those obtained using conventional micropipette aspiration.

**Figure 4:**
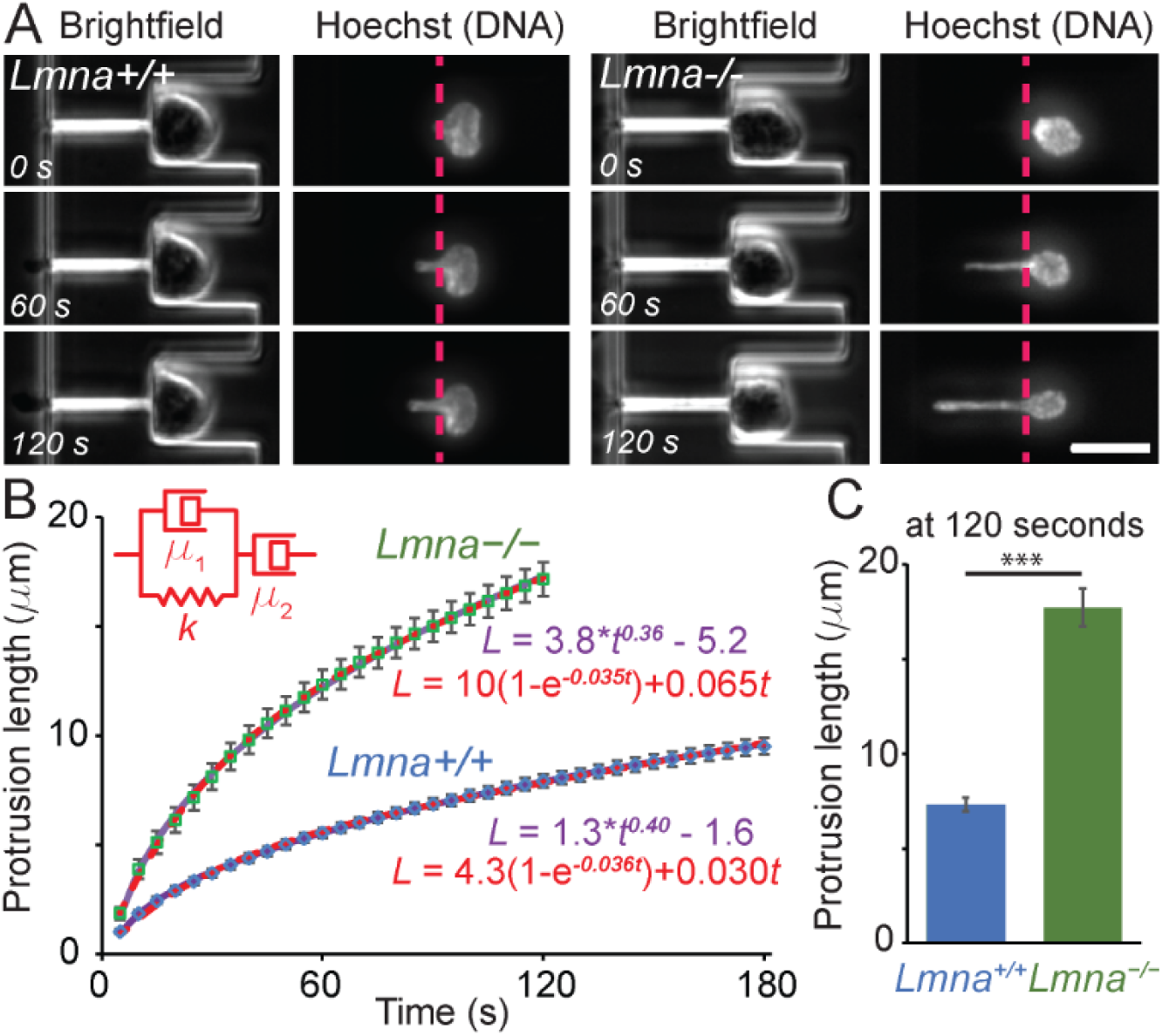
Validation of the devices the mechanical properties of nuclei. Wild-type (*Lmna*+/+, left) and lamin A/C-deficient (*Lmna*–/–, right) cells were deformed and the length of the protrusion was measured as a function of time. Brightfield images and images of the nucleus stain (Hoechst 33342) were acquired every five seconds. The *Lmna*–/– cells deformed more rapidly and more extensively than the wild-type controls. (**A**) Representative example images of the same cell at three different time points. (Scale bar 20 µm.) (**B**) The nuclear deformation (protrusion length) as a function of time modelled as a power law (purple line) or using the Jeffreys model (red dashed line). Only the first 120 seconds are shown for the *Lmna*–/– cells as many of these nuclei completely entered the micropipette channel at times longer than 120 seconds, and could thus not be used for analysis. (**C**) Comparison of the nuclear protrusion length at 120 seconds. ***, *p* < 0.001; *n* = 70 and 56 for *Lmna+*/+ and *Lmna*–/–, respectively.

In a second approach, we used classical “spring and dashpot” viscoelastic models to describe the time-dependent nuclear deformation into the micropipette channels. We tested several combinations of springs and dashpots (see Suppl. Fig. 3). The simplest model to adequately fit the observed viscoelastic creep behavior (with an increasing plateau at long deformation times) is a dashpot in series with a Kelvin-Voigt element (spring and dashpot in parallel, Figure 4B). This 3-element model, known as a Jeffreys model, predicts the time-dependent deformation by the following equation:

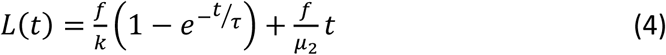

where *L*(*t*) is the strain (or, in this case, the nuclear protrusion), *f* is the aspiration force, *k* is the spring constant, µ is the dissipation coefficient of the dashpot element in series and ⍰ is the relaxation time (equivalent to k/µ_1_). To obtain quantitative data from this model, we balanced the aspiration force with the forces due to the elastic contribution (at short time scales) and the viscous flow through a small constriction (at long time scales) and obtained the following equation (see Supplementary Information for details on the derivation):

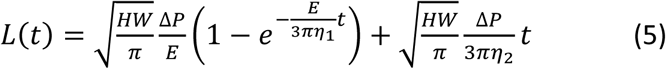

Fitting the experimental data to the Jeffreys model we obtained values comparable to those reported previously in the literature (Table 1). Guilak et al.^49^ measured an elastic modulus of 1 kPa and a viscosity of 5 kPa*s in isolated nuclei of pig chondrocytes, Dahl et al.^25^ measured an elastic modulus of 5.7 kPa in isolated nuclei from lung epithelial cells, and Luo et al^50^ found elastic moduli of 3.5 and 3 kPa in whole cell measurements of two tumor cell lines in microfluidic devices.

**Table 1.** Lamin A/C-deficient cells have altered nuclear viscoelastic properties. Parameters for the Jeffreys model based on the least squares regression of the experimental data. The parameters A and α were obtained by measuring the protrusion length in µm and the time in seconds. The units of the parameter A are dependent on the magnitude of *α*; *α* is dimensionless.

| Parameter | *Lmna*<sup>+/+</sup> | *Lmna*<sup>-/-</sup> |
| --- | --- | --- |
| A | 1.34 (±0.05) | 3.8 (±0.3) |
| α | 0.41 (±0.01) | 0.37 (±0.01) |
| E | 2.1 (±0.3) kPa | 1.0 (±0.1) kPa |
| η<sub>1</sub> | 6 (±1) kPa*s | 2.8 (±0.4) kPa*s |
| η<sub>2</sub> | 33 (±4) kPa*s | 15 (±2) kPa*s |

As expected, we detected significant differences between the lamin A/C-deficient and wild-type cells (Table 1). The elastic modulus of wild-type nuclei was more than two times larger for the lamin A/C-deficient nuclei, indicative of the importance of lamin A/C in determining the resistance to nuclear deformation.^1,4,13,17^ Similarly, the two parameters describing the nuclear viscosity were approximately double in magnitude for wild-type cells compared to the lamin A/C-deficient cells, indicating that wild-type nuclei flow more slowly.

Both the Jeffreys model and the power law model closely matched the experimental data (Figure 4B) and present complementary approaches to analyze nuclear deformation data. Taken together, the above experiments demonstrate that the microfluidic device is well suited to study nuclear mechanical properties, including the time-dependent behavior of nuclear deformation under force. Given the similar quality of fit and the fact that both viscoelastic models use the same number of tunable parameters (*A*, ⍰, and *c* for the power law model; *E, η*_1_, and *η*_2_ for the Jeffreys model), the choice of a particular model will depend on the specific experiments and questions.

### Measurements are independent of nuclear size or DNA labeling

To test the robustness of the microfluidic analysis platform in measuring nuclear mechanical properties, we analyzed the effect of two potentially confounding factors: (1) nuclear size; (2) the Hoechst 33342 dye commonly used to fluorescently label DNA, which could potentially affect nuclear deformability as it intercalates into the DNA. We found no significant correlation between the measured mechanical properties of the nuclei and the size of the nuclei (Suppl. Fig. 2C), indicating that the obtained measurements are independent of nuclear size. Furthermore, the addition of Hoechst 33342 dye did not alter the nuclear mechanical properties of cells expressing histone H2B fused to mNeonGreen to visualize nuclear deformation (Suppl. Fig. 2B), indicating that the DNA-intercalating dye does not alter mechanical properties under the experimental conditions used here.

### Application of the device to laminopathy cells, stem cells, and tumor cells

To demonstrate the versatility of the microfluidic devices in a broad range of applications, we performed measurements of nuclear mechanical properties in a variety of cell types. In the first application, we compared human skin fibroblasts from an individual with dilated cardiomyopathy caused by a mutation in the *LMNA* gene (*LMNA*-DCM) with matching skin fibroblasts from a healthy family member.^42^ *LMNA* mutations lead to a wide family of diseases, collectively referred to as laminopathies, that include *LMNA*-DCM, Emery-Dreifuss muscular dystrophy (EDMD), congenital muscular dystrophy, and limb-girdle muscular dystrophy.^15^ One hypothesis to explain the often muscle-specific phenotypes in laminopathies is that the mutations affect the mechanical properties of the nucleus, rendering it less stable, and thus resulting in increased cell death in mechanically stressed tissues such as skeletal and cardiac muscle.^15^ Supporting this hypothesis, fibroblasts expressing *LMNA* mutations associated with EDMD have more deformable nuclei than cells from healthy controls in membrane stretching assays.^20^ Applying our microfluidic platform to skin fibroblasts from a laminopathy patient with *LMNA*-DCM and from a healthy family member, we found that the *LMNA*-DCM skin fibroblasts had significantly more deformable nuclei than the healthy controls (Figure 5A Table 2), indicating that the *LMNA* mutation reduces the mechanical stability of the nucleus in the *LMNA*-DCM cells. Analysis of the time-dependent creep deformation revealed that the nuclei of the *LMNA*-DCM fibroblasts were less viscous than the healthy controls, as visible in the steeper slope of the nuclear protrusion over longer time scales (Figure 5A; Table 2). This trend recapitulates our above findings in the lamin A/C-deficient and wild-type MEFs, where the loss of lamin A/C reduced the nuclear elastic modulus and viscosities (Table 1). While further studies will be necessary to determine if these phenotypes are recapitulated in other mutations and in *LMNA* mutant human cardiomyocytes, we have already used the microfluidic assay to demonstrate that myoblasts from mouse models of muscle laminopathies have reduced nuclear stability, and that the extent of the defect correlates with the disease severity.^51^

**Table 2.**
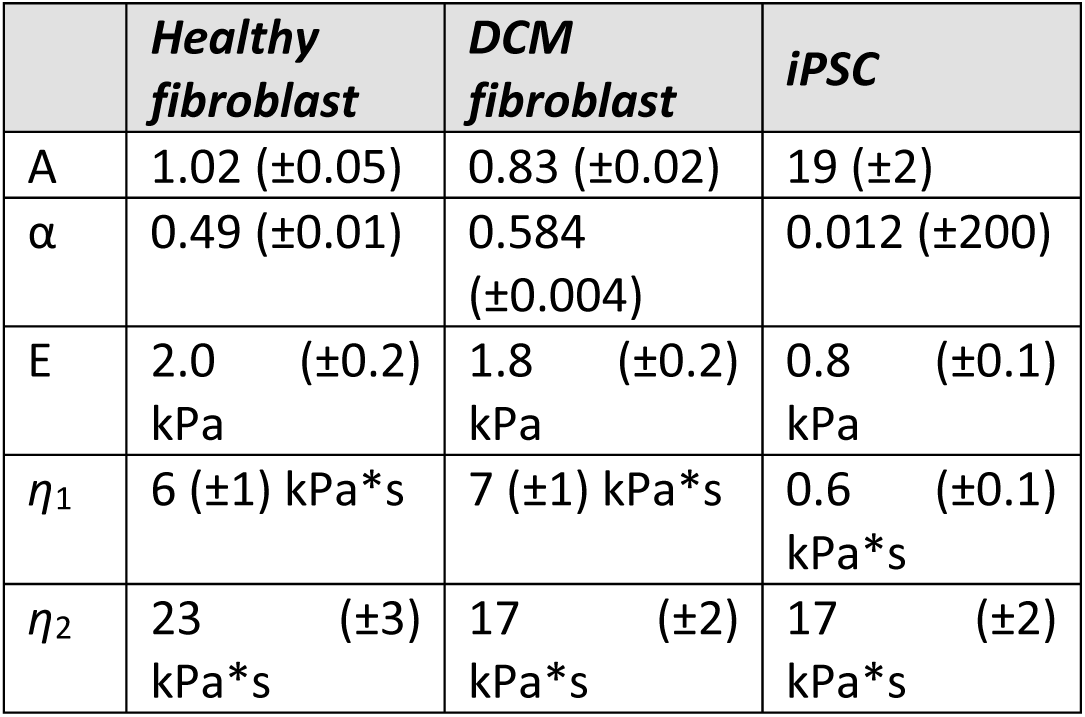
Cells bearing *LMNA* mutations and in a reprogrammed differentiation state show altered nuclear mechanics. Parameters for Jeffreys model based on best fit to the experimental data from human skin fibroblasts from an individual with an *LMNA* mutation associated with dilated cardiomyopathy (DCM), a healthy control, and iPSCs derived from healthy human skin fibroblasts. The parameters A and α were obtained by measuring the protrusion length in µm and the time in seconds. The units of the parameter A are dependent on the magnitude of α; α is dimensionless.

**Figure 5.**
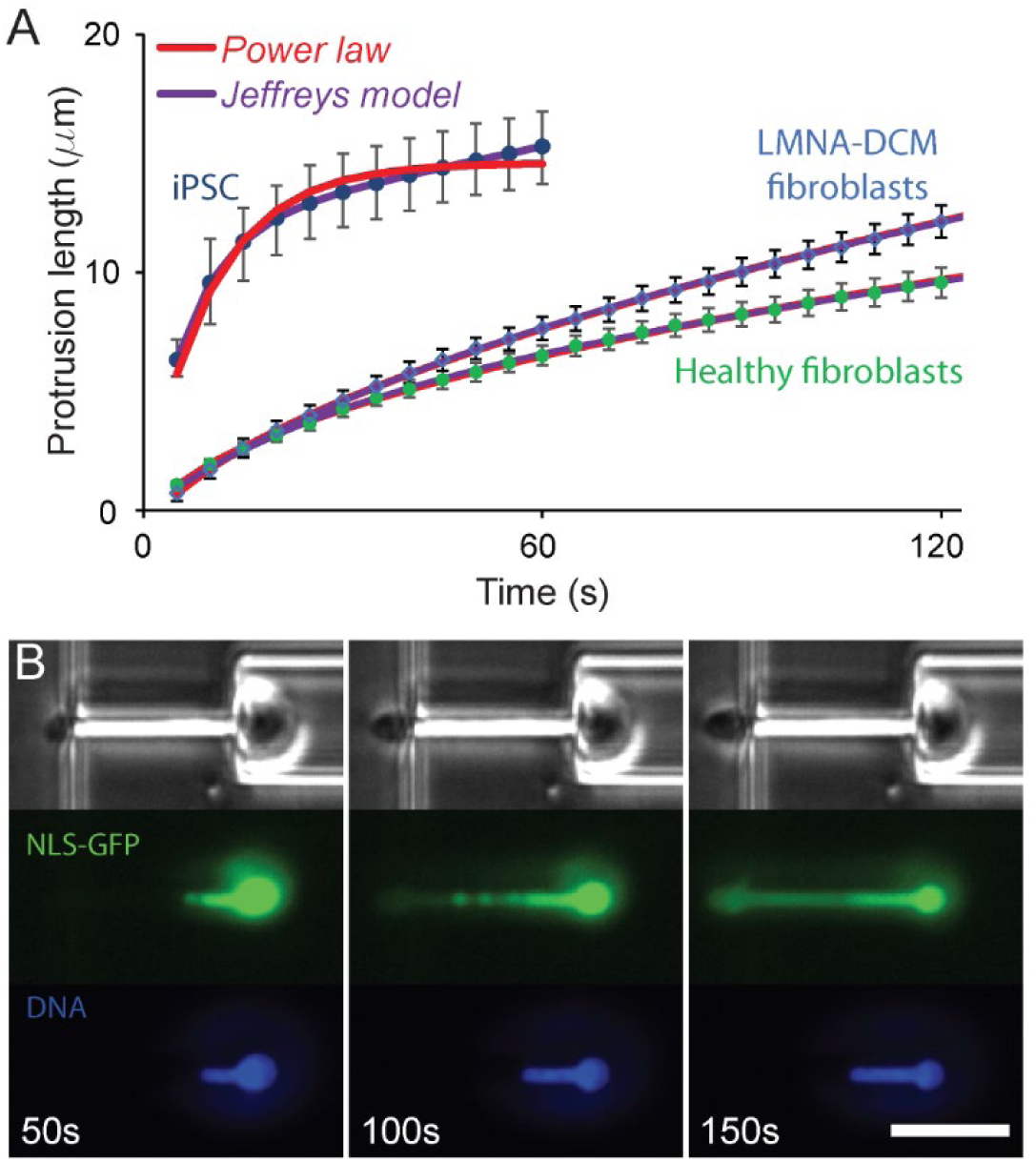
Comparison of the deformability of human cells. (**A**) Induced pluripotent stem cells derived from human skin fibroblasts have more deformable nuclei than human skin fibroblasts, reflecting the changes in chromatin organization and lower lamin A/C levels in the iPSCs. Human skin fibroblasts from an individual carrying a *LMNA* mutation that causes dilated cardiomyopathy have significantly more nuclear viscous flow at long deformation times. (**B**) Extensive nuclear deformation micropipette aspiration can result in nuclear envelope rupture, as visualized by the leakage of soluble green fluorescent proteins with a nuclear localization sequence (NLS-GFP) into the cytoplasm following nuclear envelope rupture. Time-lapse images show the extent of deformation and nuclear leakage with time, as a function of the onset of nuclear deformation. (Scale bar: 20 µm.)

In a second application, we investigated the effect of stem cell differentiation on nuclear mechanical properties. As pluripotent stem cells differentiate into specific lineages, their nuclear stiffness increases for most lineages, likely due to a concomitant increase in the expression levels of lamin A/C and changes in chromatin organization.^10,22^ We compared the deformability of human skin fibroblasts and induced-pluripotent stem cells (iPSCs) generated from skin fibroblasts, using our microfluidic devices. The iPSC cells had highly deformable nuclei (Figure 5A), resulting in many of the iPSCs passing through the micropipette channels within a few frames (less than 20 seconds). To avoid bias towards cells that passed through the channel more slowly, we restricted our comparison to the first 60 seconds of nuclear deformation and selected only cells whose nuclei had not completely entered the micropipette channel during time. The iPSCs had significantly more deformable nuclei than the skin fibroblasts (Figure 5A; Table 2), consistent with a previous study using conventional micropipette aspiration that found that nuclear stiffness increased during differentiation of human embryonic stem cells^22,52^. Comparing the data to both the power law model and the Jeffreys model, we found that the Jeffreys model provided a better fit for the iPSC data than the power law model, whereas both models provided equally good fits for the human skin fibroblast data (Figure 5A), consistent with our results for mouse embryo fibroblasts (Figure 4). The error on the power law exponent value is orders of magnitude greater than the exponent itself, symptomatic of the poor fit. Strikingly, the viscosity (*η*_*2*_) of the iPSCs did not differ from the viscosity of the skin fibroblasts. This viscosity governs the deformation rate at long time scales. Our results suggest that reprogramming primarily alters the elastic properties of the nuclei.

Taken together, these examples demonstrate the use of the microfluidic device to measure the viscoelastic properties of nuclei in intact cells in a broad range of applications, producing results consistent with conventional micropipette aspiration assays or nuclear strain experiments, but at significantly higher throughput, and without the need for cell-substrate adhesion. The latter point is particularly relevant when studying tumour cells, which often have reduced adhesion strength,^53^ and are thus not well suited for substrate strain experiments. Taking advantage of the novel microfluidic assay, we recently demonstrated that TGF-beta induced epithelial-to-mesenchymal transition in PyMT mouse breast tumour cells was associated with a decrease in nuclear stiffness, which, together with changes in focal adhesion organization, resulted in increased tumour cell invasion.^54^ Notably, the device can also be used to study nuclear envelope rupture, which frequently occurs during migration of cells through confined environments.^26,41,55^ As demonstrated in Figure 5B, the leakage of soluble green fluorescent protein with a nuclear localization sequence (NLS-GFP)^41^ from the nucleus into the cytoplasm upon nuclear envelope rupture can be clearly observed during large nuclear deformations.

## Conclusions

We developed a novel microfluidic device and semi-automated imaging analysis pipeline in which we can observe and quantify the deformation of the nucleus at high resolution in intact cells, and with substantially higher throughput than conventional single cell micropipette aspiration experiments or atomic force microscopy measurements. We demonstrated the device’s applicability to obtain precise viscoelastic information about the nucleus, including in mouse and human laminopathy cells and in human induced pluripotent stem cells and the corresponding original skin fibroblasts. Because the analysis platform presented here can perform measurements on large populations of cells, it can characterize the heterogeneity of samples, for example, to detect small mechanically distinct subpopulations of cancer cells or stem cells.

## Acknowledgements

This work was supported by awards from the National Institutes of Health [R01 HL082792 and U54 CA210184 to JL], the Department of Defense Breast Cancer Research Program [Breakthrough Award BC150580 to JL], the National Science Foundation [CBET-1254846 and MCB-1715606 to JL; GRFP-2014163403 to GRF], La Ligue contre le cancer [REMX17751 to PMD] the Fondation ARC [PDF20161205227 to PMD], and a ministerial doctoral fellowship of Paris Saclay University to SDM. This work was performed in part at Cornell NanoScale Facility (CNF), an NNCI member supported by NSF Grant NNCI-1542081. The authors thank Elisa di Pasquale and Gianluigi Condorelli for the human iPSCs and laminopathy patient fibroblasts, Colin Stewart for the lamin A/C-deficient and wild-type MEFs. The authors thank Karine Guevorkian and Francoise Brochard for helpful discussions.

