## Supplementary Materials & Figures for "High-throughput microfluidic micropipette aspiration device to probe time-scale dependent nuclear mechanics in intact cells"

### Supplemental Information

#### Derivations

We derived the elastic modulus and the viscosities of the viscoelastic model as laid out in previous work,[56] with adaptations for a rectangular channel and our chosen model. The Jeffreys model consists of three elements: a dashpot in series with a Kelvin-Voigt element, i.e., a spring and a dashpot in parallel. For this model, the creep is described by the following equation:

$$L(t) = \frac{f}{k}\left(1 - e^{-t/\tau}\right) + \frac{f}{\mu}t$$

Where *f* is the force, $k$ is the spring constant, $\tau$ is the relaxation time ($\tau = \frac{k}{\mu}$) and *μ* is the viscosity of the dashpot in series. At short time scales, the first term dominates, resulting in a rapidly rising curve, while at long time scales, the second term dominates, leading to a linear regime.

The aspiration force is given by:

$$f = \pi R_p^2 \Delta P$$

Where $R_p$ is the radius of the micropipette and $\Delta P$ is the applied pressure.

At short time scales, the force is balanced by the elastic deformation and is given by the following equation:

$$\frac{f}{A} = CE\frac{\delta}{R_p}$$

Where *A* is the cross-sectional area of the pipette, *C* is approximately equivalent to 1,[56] *E* is the elastic modulus and *δ* is the elastic deformation at short times. We thus obtain $f = \pi R_p E\delta$.

The definition of the spring constant (*k*) is the relationship between the force and the extension: $f = k \times \delta$, and thus $\frac{f}{\delta} = k = \pi R_p E$.

At long time scales, the force is balanced by viscous flow. The dissipative force due to the plug at the entrance of the capillary is given by the following equation:

$$f = 3\pi^2 \eta R_p \frac{dL}{dt}$$

Where *μ* is the viscosity and $R_p$ is the radius of the pipette. At long time scales, the second term of equation 1 dominates and we obtain: $\frac{dL}{dt} = \frac{f}{\mu}$, thus $\mu = \frac{f}{dL/dt} = 3\pi^2 \eta R_p$

Inserting these expressions into equation 1, we obtain the following:

$$L(t) = \frac{f}{k}\left(1 - e^{-t/\tau}\right) + \frac{f}{\mu}t = \frac{\pi R_p^2 \Delta P}{\pi R_p E}\left(1 - e^{-\frac{E}{3\pi\eta}t}\right) + \frac{\pi R_p^2 \Delta P}{3\pi^2 \eta R_p}t = \frac{R_p \Delta P}{E}\left(1 - e^{-\frac{E}{3\pi\eta}t}\right) + \frac{R_p \Delta P}{3\pi\eta}t$$

Here we replace the radius of the pipette by the effective radius for a rectangular channel, given by $R_{eff} = \sqrt{\frac{HW}{\pi}}$, Where H is the height and W is the width of the channel.[57]

We thus obtain:

$$L(t) = \sqrt{\frac{HW}{\pi}\frac{\Delta P}{E}\left(1 - e^{-\frac{E}{3\pi\eta}t}\right)} + \sqrt{\frac{HW}{\pi}\frac{\Delta P}{3\pi\eta}t}$$

By fitting the data, we can obtain values for the elastic modulus and the viscosities.

### Supplementary Figures

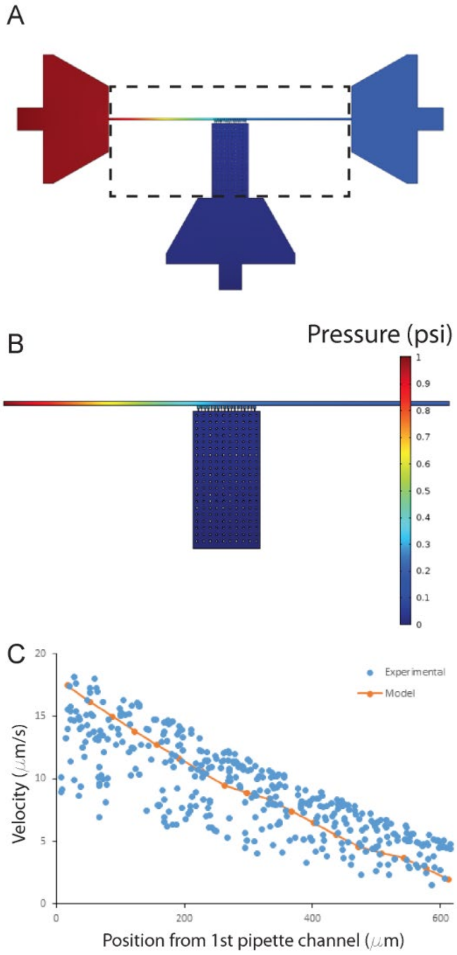


**Supplementary Figure 1: Simulation on micropipette channels in "open" configuration.** (**A**) Overview of the simulated pressure distribution in the entire microfluidic device. (**B**) Close-up of the area containing the micropipette channels (area surrounded by dashed line in panel A) showing a rapid decrease of the pressure upstream of the micropipette channels. (**C**) Comparison of the velocity obtained from the simulation (orange line) and measured velocities obtained from the streak length of fluorescent beads in the microfluidic device (blue points).

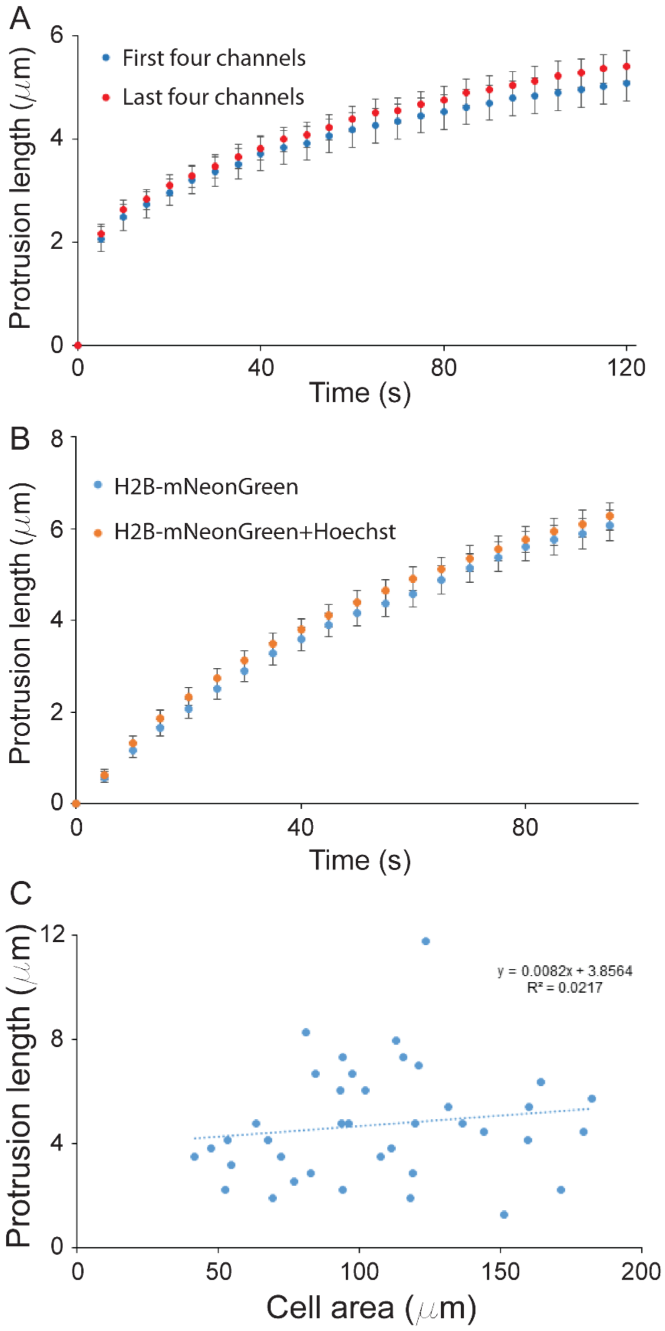


**Supplementary Figure 2. Effect of microchannel position, Hoechst labeling, and cell size on nuclear deformation measurements. (A)** The deformation rate as a function of microchannel position. We compared the deformation rate or MDA-MB-231 cells in the first four and last four micropipette channels. Similar to the results shown in Figure 3C, the position of the cells does not have a significant influence on the deformation rate of the cells. **(B)** Effect of a DNA intercalating agent on the deformability measurements. The deformability of cells was not significantly different when Hoechst 33342 was added at the concentrations we use in our experiments. (**C**) Effect of nuclear size on deformability of the nucleus. We compared the cross-sectional area and the protrusion length at 60 s in individual cells. Using a regression analysis, we determined that the slope of the linear regression (0.008) was smaller than the 95% confidence interval (0.018) associated with it, indicating that the slope is not significantly different from 0.

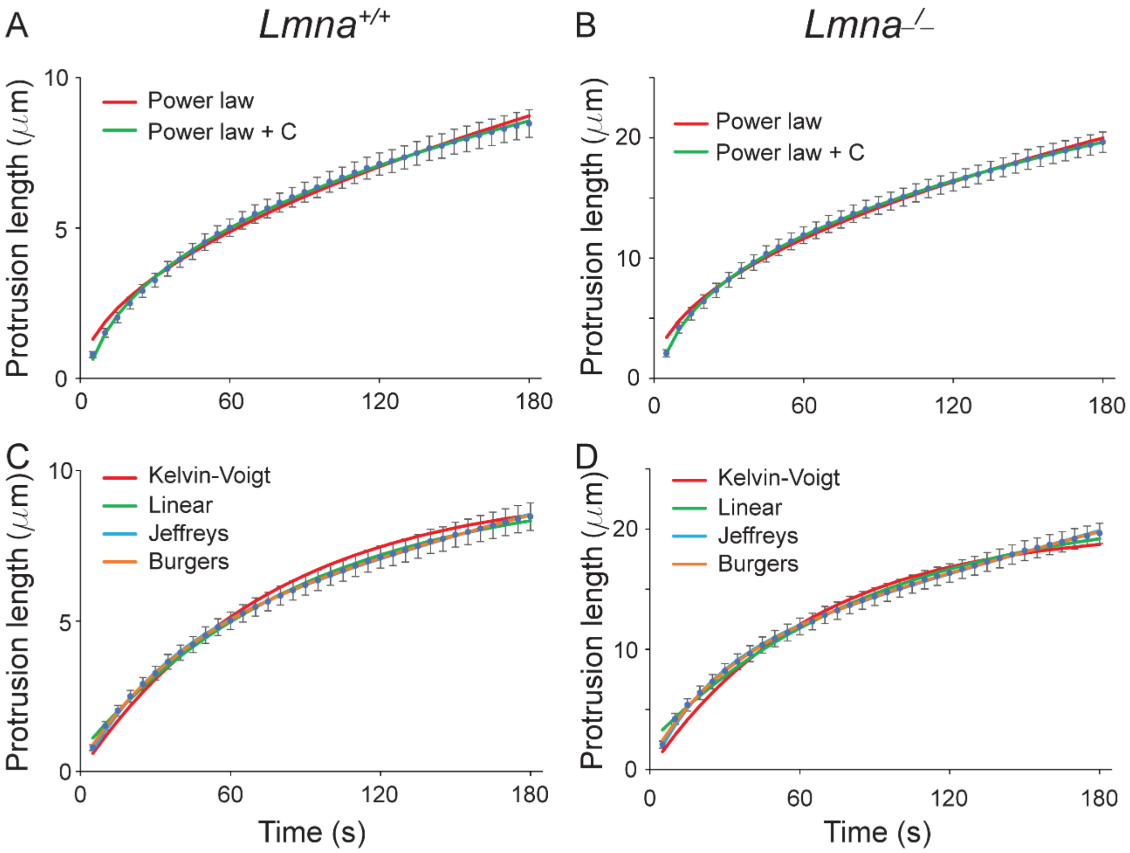


**Supplementary Figure 3. Comparison of the models used to fit the data.** (**A, B**) The data was fit with a simple power law ($y=At^{\alpha}$) and a power law with an additional constant ($y=At^{\alpha}+C$) to account for error in the first time point. The simpler model does not follow the data well at small time points due to the uncertainty in time zero, resulting in a flattening of the curve and thus a large change in the exponent value. The second model is a better fit at short time points. (**C, D**) The four viscoelastic (spring and dashpot) models. The Kelvin-Voigt and Standard Linear Solid models include a spring in parallel and thus do not result in a viscous linear increase with time. We thus didn't chose these models as our data increases linearly at long time points (indicative of a dashpot in series). The Jeffreys model is made up of a dashpot in series with a Kelvin-Voigt element and thus is the simplest element to accurately describe our data. Accordingly, the $R^2$ values are smaller for the Jeffreys model than the first two models. (See Suppl. Table 1.) The Burgers model includes an additional element, and consequently exhibits a better fit, but is only a minimal improvement.

**Supplementary Tables**

| ***Model*** | ***# of Variables*** | ***Lmna***$^{+/+}$ | | ***Lmna***$^{-/-}$ | |
|---|---|---|---|---|---|
| | | **$R^2$** | **Residuals** | **$R^2$** | **Residuals** |
| Power law | 2 | 0.99345 | 1.1 | 0.99520 | 3.7 |
| Power law with C | 3 | 0.99941 | 0.11 | 0.99989 | 0.13 |
| Kelvin-Voigt | 2 | 0.99401 | 2.4 | 0.98903 | 13 |
| Linear | 3 | 0.99800 | 0.32 | 0.99639 | 3.9 |
| Jeffreys | 3 | 0.99963 | 0.082 | 0.99921 | 0.68 |
| Burgers | 4 | 0.99980 | 0.031 | 0.99941 | 0.37 |

**Supplementary Table 1: Values of the Coefficient of Determination ($R^2$) and residuals calculated for each condition**
